# Bioinformatics analysis identifies sequence determinants of enzymatic activity for the PHARC associated lipase ABHD12

**DOI:** 10.1101/2025.02.19.639194

**Authors:** Arnab Chakraborty, Archit Devarajan, Kundan Kumar, C. S. Rohith, M. S. Madhusudhan, Girish S. Ratnaparkhi, Siddhesh S. Kamat

## Abstract

In humans, PHARC (***<u>p</u>***olyneuropathy, ***<u>h</u>***earing loss, ***<u>a</u>***taxia, ***r***etinitis pigmentosa, and ***<u>c</u>***ataract) is an early onset autosomal recessive neurological disorder caused by deleterious mutations to ABHD12 (α/β-hydrolase domain protein # 12). Biochemically, ABHD12 functions as a lipase, and catalyzes the hydrolysis of lysophosphatidylserine (lyso-PS) (lyso-PS lipase). By doing so, it controls the concentrations and signaling pathways regulated by this potent signaling lysophospholipid in the mammalian brain. While genetic mapping efforts have identified over 30 mutations in ABHD12 from human PHARC subjects, the biochemical activity of these pathogenic mutants remains unknown. To understand this, here, we performed an exhaustive bioinformatics survey, and collated ABHD12 protein sequences from various organisms across evolution. Next, based on sequence alignments and structural modeling, we identified functionally relevant conserved residues in the ABHD12 protein sequence that are potentially important for its enzymatic activity. To validate these *in silico* findings, we generated numerous mutants of murine ABHD12, including those associated with human PHARC subjects, and assayed them for their enzymatic activity. Taken together, these complementary *in silico* and biochemical studies provide the first thorough sequence-function relationship for mammalian ABHD12, especially relevant in the context of PHARC. Finally, our evolutionary analysis identified CG15111 as an ABHD12 ortholog in the fruit fly (*Drosophila melanogaster*), and enzymatic assays indeed confirmed that recombinant CG15111 has robust lyso-PS lipase activity. Flies serve as an excellent animal system to model various human neurological diseases, and the identification of CG15111 as a *Drosophila melanogaster* ABHD12 ortholog opens new avenues to study PHARC in fly models.

## INTRODUCTION

Over a decade ago, an autosomal recessive genetic disorder named PHARC (acronym for all the symptoms associated with the disease: ***<u>p</u>***olyneuropathy, ***<u>h</u>***earing loss, ***<u>a</u>***taxia, ***r***etinitis pigmentosa, and ***<u>c</u>***ataract; OMIM: 612674) was reported in humans^1–5^. Clinically, human PHARC subjects show early onset visual disturbances (e.g. cataract or partial blindness) and auditory deficits (e.g. deafness), that are often coupled to polymodal sensory and motor defects caused by peripheral neuropathy (e.g. *pes cavus*)^1, 6, 7^. The symptoms associated with PHARC in human subjects manifest in early teenage years, progressively worsen with age, and there seems to be no gender or race bias associated with this genetic disease^6, 7^. In extreme cases, demyelination of sensorimotor neurons and cerebellar atrophy has also been reported in older human PHARC subjects^1, 6, 7^, and yet, to date, there are no reported medical cures or therapies to treat this human genetic neurological disorder. Given the array of symptoms, PHARC was commonly misdiagnosed as possibly other neurological disorders, until genetic mapping confirmed that this neurodegenerative disease was caused by deleterious mutations in the *abhd12* gene on chromosome 20p11 in humans^1, 8^. To date, over 30 pathogenic mutations have been reported in the *abhd12* gene, all causing PHARC in humans^6, 7^. These mutations include deletion-insertion, nonsense, frameshift, splice site, and missense type mutations in the *abhd12* gene from samples that were genetically sequenced from human PHARC subjects^6^.

The *abhd12* gene encodes an integral membrane associated lipase, α/β-hydrolase domain containing protein # 12 (ABHD12), which belongs to the metabolic serine hydrolase family of enzymes^9, 10^. Untargeted lipidomics analysis on the brains of ABHD12 knockout mice (the murine model of PHARC) showed that relative to wild-type controls, the loss of ABHD12 activity resulted in a massive accumulation of different lysophosphatidylserine (lyso-PS) lipids^11^. Lyso-PSs are a class of signaling lysophospholipids, that have many important functions in mammalian physiology, and dysregulation in their metabolism is now causally linked to an array of human neurological and autoimmune disorders, including PHARC^12, 13^. From a biochemical standpoint, mechanistic studies have since shown that via its enzymatic activity, ABHD12 catalyzes the degradation of lyso-PS lipids, and, by doing so, functions as the principal lyso-PS lipase (**Figure 1**) in the mammalian central nervous and immune systems^11, 14^.

**Figure 1.**
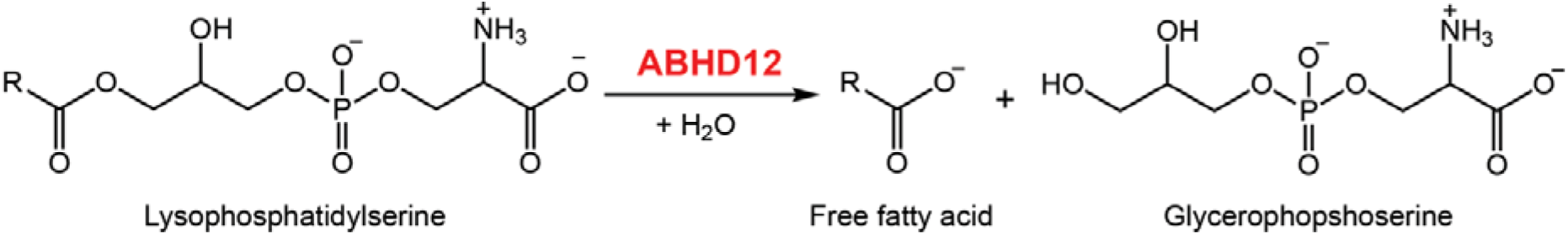
The lyso-PS lipase reaction catalyzed by ABHD12. In this reaction, R denotes long or very-long chain fatty acids.

Following up on these studies, we have shown that ABHD12 is predominantly localized to the endoplasmic reticulum (ER) membrane, and has a luminal orientation of the enzyme active site, that regulates the secretion of lyso-PS lipids from different mammalian cells^15, 16^. Further, from *in vitro* biochemical assays, we demonstrate that ABHD12 prefers very-long chain (VLC) variants of lyso-PS as substrates^16, 17^. This substrate preference of ABHD12 explains why its deletion results in the highest accumulation of pro-inflammatory VLC lyso-PSs in the mammalian brain, that are largely responsible for neuroinflammation and eventually the neurobehavioral symptoms associated with human PHARC subjects^17, 18^. Finally, by immunohistochemical analysis coupled with biochemical assays and targeted lipidomics measurements, we have reported that ABHD12 has ubiquitous expression and activity in different anatomical regions of the mammalian brain, and the loss of ABHD12 activity results in dysregulated lyso-PS metabolism in all anatomical regions of the mammalian brain without exception^19^. These studies from our lab, together with previous literature, provide a nice explanation for the progression of PHARC^6, 7^, and ABHD12’s role in the pathogenesis of this neurodegenerative disease.

While significant studies have been done to delineate how the biochemical activity of ABHD12 regulates lyso-PS metabolism and, the association of the ABHD12-lyso-PS signaling axis during PHARC, the absence of any experimentally solved three-dimensional structure has resulted in only a limited understanding of the protein sequence-activity relationship of ABHD12. This can be particularly important in the context of PHARC, as newer ABHD12 mutants are mapped in humans suffering from neurological symptoms via genetic sequencing efforts. To address this knowledge gap, in the study, we perform a thorough bioinformatics survey for the identification of ABHD12 protein sequences across all forms of life. Next, we phylogenetically classify these putative ABHD12 sequences across the evolutionary time scale and find positionally and functionally conserved residues in their protein sequences. Further, using structural modeling and docking studies, we identify residues that might have importance in the biochemical activity of ABHD12 and, coupled with biochemical assays, show that these residues are indeed of importance for the enzymatic activity of ABHD12. We also find that several missense mutations associated with PHARC, occur in highly conserved residues, and these mutants are catalytically compromised. Finally, from our evolutionary analysis, we find that ABHD12 has an ortholog in flies, CG15111, that also robustly performs lyso-PS lipase activity, and thereby opens new prospects in combination with genetics for modeling pathogenic mutants and thus studying PHARC in fly models.

## MATERIALS AND METHODS

### Materials

Unless otherwise mentioned: all chemicals, buffers, and reagents were procured from Sigma-Aldrich; all mammalian tissue culture media and consumables were procured from HiMedia Laboratories; all LC-MS grade solvents were procured from JT Baker. Wherever applicable, the catalog numbers of materials used in this study are mentioned below. All oligonucleotides used in this study were obtained from Sigma-Aldrich.

### Bioinformatics analysis

The bioinformatics analyses were performed as previously reported by us with modifications^20^. Briefly, five iterations of PSI-BLAST searches on the human ABHD12 (hABHD12) (UniProt: Q8N2K0, NCBI: NP_001035937.1) as the reference sequence against the clustered non-redundant (clustered_nr) database was performed to identify distantly-related orthologs of ABHD12^21^. The expect threshold (E-value) was set to 1× 10^−6^ with a maximum return of 1000 hits. The individual members of each cluster (totaling to 2770) were extracted and subject to further analysis. A PSI-BLAST hit was classified as ABHD12 protein sequence if it met all the following criteria: (i) has an N-terminal transmembrane domain, (ii) conserved catalytic triad, (iii) conserved nucleophilic (YIW<u>G</u>H<u>S</u>L<u>G</u>TGV) and acyltransferase (LGY<u>H</u>VVTF<u>D</u>YRG) motifs with up to three mismatches in residues other than the underlined residues (iv) sequence length between 250 and 550 amino acids. DeepTMHMM^22^ and CCTOP^23^ were used to assess the presence of any transmembrane domains. A previously reported custom script was slightly modified and used to further curate the data based on the presence of the catalytic triad and other key functional residues by carrying out pairwise global alignments using the Needleman-Wunsch algorithm^24^. For plotting the phylogenetic tree, the longest isoform of ABHD12 from each organism was selected and subject to multiple sequence alignment using Clustal Omega^25^. The resulting alignment was fed into MEGA11^26^ to plot maximum likelihood trees using the James-Taylor-Thornton (JTT) model^27^ and the trees were visualized in iTol^28^. The highly conserved ABHD12 residues were identified using pymsaviz (pypi.org/project/pymsaviz/) and Bio.Align (biopython.org/docs/1.76/api/Bio.Align.html) packages in Python. The functionally conserved residues were identified by manual inspection of filtered protein sequences.

### Structural docking studies

All structural docking studies were performed on the AlphaFold^29, 30^ generated structure of hABHD12 (UniProt ID: Q8N2K0). To identify a substrate binding pocket that contained the catalytic serine (Ser-246) within hABHD12, a tunnel analysis was performed using the CAVER Web 1.0^31^ and MOLE 2.5^32^ online servers. Following this, surface electrostatic calculations were performed on the same AlphaFold structure of hABHD12 using the APBS plugin to PyMOL (https://pymolwiki.org/index.php/Apbsplugin) to understand the charge distribution on the protein surface and in the active site. The PARSE force field was used to generate the required PQR files which were subsequently used to calculate the electrostatic surface diagrams. Finally, ligand (C18:0 lyso-PS) docking was performed on the same AlphaFold structure of hABHD12 using Autodock Vina in the PyRx environment using standard reported procedures. Briefly, the structure of hABHD12 and the SDF files of the ligand (C18:0 lyso-PS) was converted into the PDBQT file format using OpenBabel^33^. Thereafter, targeted docking was performed with a (30 x 30 x 30) A box roughly centered at the catalytic serine (Ser-246) in the identified ligand binding pocket of hABHD12, and the top nine binding conformations were selected for further investigation.

### Overexpression of mouse ABHD12 mutants in HEK293T cells

The *abhd12* gene (encoding ABHD12) was amplified from an in-house generated cDNA library of the mouse brain, and cloned into a *pCMV-Sport6* vector as reported previously^16^. All mutants of mouse ABHD12 (mABHD12) were made in the same plasmid using standard site-directed mutagenesis protocols as per manufacturer’s instructions (Promega). All primers used to generate the mABHD12 mutants can be found in **Table S1**. Wild type (WT) or mutants of mABHD12 were transiently transfected into mammalian HEK293T cells by an established methodology using the polyethylenimine-based PEI MAX® transfection reagent^15–17^. As a control in these experiments, an empty plasmid (mock control) was used to account for off-targets (if any) from the empty plasmid during these studies. Two days post-transfection, the cells were harvested by scraping, washed with cold sterile Dulbecco’s phosphate buffered saline (DPBS) (3 times), and then lysed by sonication. The cellular debris (pellet) was discarded by centrifugation of the lysate at 200g for 5 min at 4 °C, and the resulting supernatant was further centrifuged at 100,000g for 45 min at 4 °C to obtain the membrane proteomic fraction as reported earlier^15–17^. The activity and overexpression of mABHD12 variants in membrane proteomic fractions of HEK293T cells were confirmed by gel-based ABPP assays and Western blot analysis respectively.

### Overexpression of CG15111 in insect cells

The *cg15111* gene (encoding CG15111) was amplified from a laboratory generated cDNA library of the adult fly, and cloned with a C-terminal FLAG tag into the *pRmHa3* vector^34^. The catalytic serine (Ser-241) was mutated to an alanine to yield the S241A CG15111 mutant in the same plasmid using standard site-directed mutagenesis methods as per manufacturer’s instructions (Promega). All primers used for the amplification and site-directed mutagenesis studies are listed in **Table S1**. The overexpression of WT or S241A CG15111 was performed in Schneider 2 (S2) insect cells using transfection protocols recently reported by us^35^. The activity and overexpression of CG15111 variants in membrane proteomic fractions of S2 cells were confirmed by gel-based ABPP assays and Western blot analysis respectively.

### Gel-based ABPP assays

All gel-based ABPP assays were done using protocols previously reported by us^15–17, 36^. Briefly, all protein concentrations were estimated using Bradford’s reagent (Sigma-Aldrich, catalog # B6916), and 50 μg of the membrane proteomic fraction (of HEK293T cells or S2 cells) was incubated with 2 μM FP-rhodamine (45 mins, 37 °C) with constant shaking. Following this, the reaction was quenched using 4x-SDS loading buffer as reported earlier^15–17, 36^. All gel-based ABPP samples were loaded on a 10% SDS-PAGE gel, and enzymatic activities (of ABHD12 or CG15111 variants) were visualized by in-gel fluorescence on an iBright1500 gel documentation system (Invitrogen).

### Western blot analysis

Following gel-based ABPP assays, the resolved membrane proteomic fractions were transferred to a nitrocellulose membrane (GE Healthcare, catalog # GE10600002) and processed for Western blot analysis using protocols previously reported by us^15, 16, 37^. In the Western blot analysis, Ponceau S staining (Sigma-Aldrich, catalog # A40000279) was used to confirm equal loading of protein samples. The primary antibodies used in our Western blot studies were: anti-ABHD12 (rabbit, Abcam, catalog # ab68949) and anti-FLAG (rabbit, Sigma-Aldrich, catalog # F7425). The secondary antibody used in our Western blot studies was an anti-rabbit conjugated to horseradish peroxidase (HRP) (Thermo Fisher Scientific, catalog # 31460). All Western blots were developed with the Immobilon chemiluminescent HRP substrate (Merck Millipore, catalog # WBKLS0500) and visualized on a G-Box Chemi-XRQ gel documentation system (Syngene).

### Lyso-PS lipase assays

All LC-MS based lyso-PS lipase assays were done by LC-MS analysis using procedures previously reported by us^15–17, 19^. All lyso-PS lipase assays were done with 20 μg of the membrane proteomic fraction (from HEK293T or S2 cells) against 100 μM lyso-PS substrate (C17:1 lyso-PS; Avanti Polar Lipids, catalog # 858141) as reported earlier^15–17, 19^. All assays were analyzed on an Agilent 6545 Quadrupole Time Of Flight (QTOF) LC-MS/MS instrument in the negative ion mode using an electrospray ionization (ESI) source. All LC separations were done using columns and solvent gradients previously reported by us^15–17, 19^. The MS parameters used were as follows: drying and sheath gas temperature = 320 °C; drying and sheath gas flow rate = 10L/min; fragmentor voltage = 150 V; capillary voltage = 4 kV; nebulizer (ion source gas) pressure = 45 Ψ and nozzle voltage = 1 kV. The product release was quantified by measuring the area under the curve for the peak corresponding to C17:1 FFA (produced from C17:1 lyso-PS), and normalizing it to the internal standard. The substrate hydrolysis rate was corrected by subtracting the non-enzymatic rate of hydrolysis, which was obtained by using corresponding heat-denatured (15 min at 95 °C) membrane proteomes.

### Data plotting and analysis

All graphs represented in this study are analyzed and plotted using the GraphPad Prism 10 (version 10.3.0) software for macOS.

## RESULTS

### Identification of ABHD12 protein sequences

ABHD12 is an integral membrane associated lipase from the metabolic SH family, with one predicted N-terminal transmembrane helix, and an invariant catalytic triad^19^. In the absence of any experimentally solved structures of ABHD12, the N-terminal transmembrane helix is predicted to comprise of residues 75 to 92, while the catalytic triad is expected to comprise of Ser-246, Asp-333 and His-372 for human ABHD12 based on a protein sequence analysis^19^ (**Figure 2**). Further, a previous bioinformatics survey of various ABHD-enzymes suggests that the conserved nucleophilic serine residue essential for catalysis (e.g. Ser-246 for human ABHD12) is part of a canonical nucleophilic GxSxG motif (where x = any amino acid)^38^ (**Figure 2**). The same survey also finds a putative acyltransferase motif comprising of a HxxxxD motif (x = any amino acid) within the conserved LGY<u>H</u>VVTF<u>D</u>YRG sequence for mammalian ABHD12 sequences^38^ (**Figure 2**). Hence, based on these bioinformatics studies, we set a stringent threshold, and classified a particular protein sequence as ABHD12 only if it possessed the N-terminal transmembrane domain (helix), conserved catalytic triad, nucleophilic motif and acyltransferase motif. For the acyltransferase motif, the His and Asp of the HxxxxD segment were set as a constant, and up to three mismatches were permitted within the remaining LGY<u>H</u>VVTF<u>D</u>YRG sequence. Finally, we restricted the protein sequence length from 250 to 550 amino acids, to ensure that any promiscuous sequences were filtered out of the analysis.

**Figure 2.**
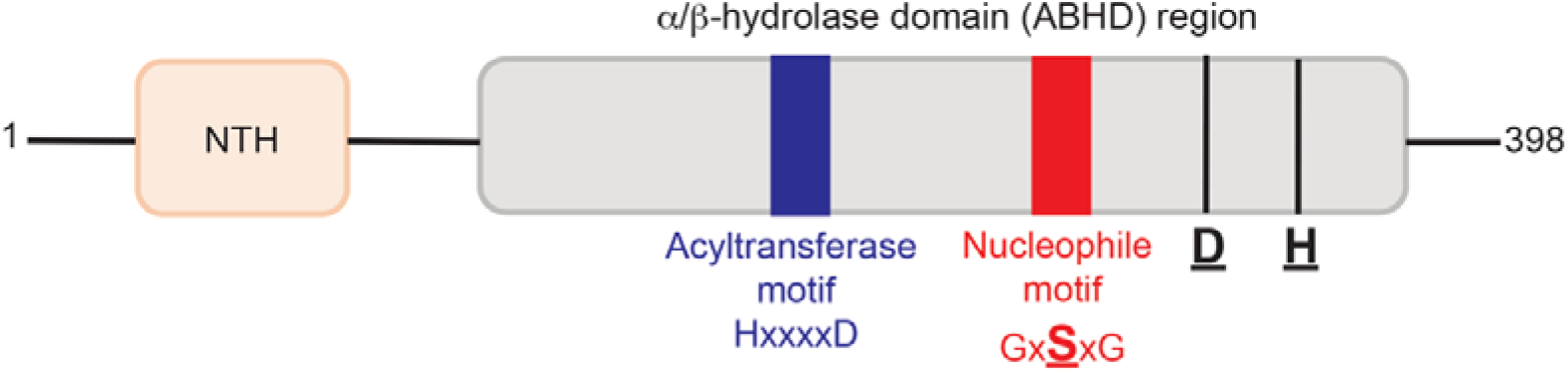
Schematic representation of the hABHD12 structure. hABHD12 is predicted to contain the canonical catalytic triad consisting of Ser-Asp-His, a nucleophilic motif that contains the serine residue as part of a GxSxG sequence, and an acyltransferase motif comprising of an invariant HxxxxD sequence within the ABHD region of the protein. Additionally, hABHD12 also contains a N-terminal transmembrane helix (NTH) that anchors this lipase to the ER membrane.

After setting appropriate selection criteria to obtain a list of possible ABHD12 protein sequences, five independent rounds of iterative PSI-BLAST searches were performed on the clustered non-redundant (clustered_nr) sequences database using human ABHD12 (UniProt: Q8N2K0, NCBI: NP_001035937.1) as the query sequence (**Table S2**). This search resulted in the identification of 2770 protein sequences that were then subjected to transmembrane domain prediction using two separate web servers for further filtering. A N-terminal integral transmembrane helix was identified in 2362 of these protein sequences, of which the transmembrane regions in 49 protein sequences were identified by only one of the two web servers. Among these, upon further filtering, only 2223 protein sequences contained the catalytic triad and satisfied the protein length criteria for further selection. After a final round of refinement, a total of 1425 protein sequences that contained both the nucleophilic and acyltransferase motifs and could be reliably classified as ABHD12 based on passing all our set criteria (**Table S2**).

### Phylogenetic analysis of ABHD12 protein sequences

An inspection of the 1425 protein sequences identified as ABHD12 from the bioinformatics analysis showed that several organisms had more than one isoform of ABHD12. Therefore, to retain maximum species-specific information for the ABHD12 protein sequences that satisfy all filtering criteria, the longest isoform from each organism (except human, wherein the query sequence was used) was manually selected. This narrowed down the list of ABHD12 protein sequences to 860 across the same number of unique organisms. Next, to understand the evolutionary time scale of ABHD12, we performed a phylogenetic analysis on these 860 ABHD12 protein sequences (**Figure 3A**). This phylogenetic classification showed that two phyla, *Chordata* and *Arthropoda*, almost exclusively contained all the ABHD12 protein sequences. Amongst them, the phylum *Chordata* had the highest prevalence of ABHD12 protein sequences, (727 of 860, 84.5%), with phylum *Arthropoda* containing most of the remaining ABHD12 protein sequences (117 of 860, 13.6%) (**Figure 3B**). Within phylum *Chordata*, ABHD12 protein sequences were most prevalent in class *Aves* (birds) (43.1%, 313 of 727 chordates), *Actinopterygii* (bony fish) (25%, 182 of 727 chordates) and *Mammalia* (mammals) (24.8%, 180 of 727 chordates), while a smaller fraction of ABHD12 protein sequences were also found in classes *Reptilia* (reptiles) (5.9%, 43 of 727 chordates) and *Amphibia* (amphibians) (1.2%, 9 of 727 chordates) (**Figure 3C**). From an evolutionary perspective within phylum *Chordata*, the phylogenetic analysis suggests that the ABHD12 protein sequences from reptiles-birds and mammals-amphibians cluster together, while those from fish form a distinct outgroup (**Figure 3A**). Notably, amongst phylum *Arthropoda*, class *Insecta* (insects) contains the highest number of ABHD12 protein sequences (81.2%, 95 of 117).

**Figure 3.**
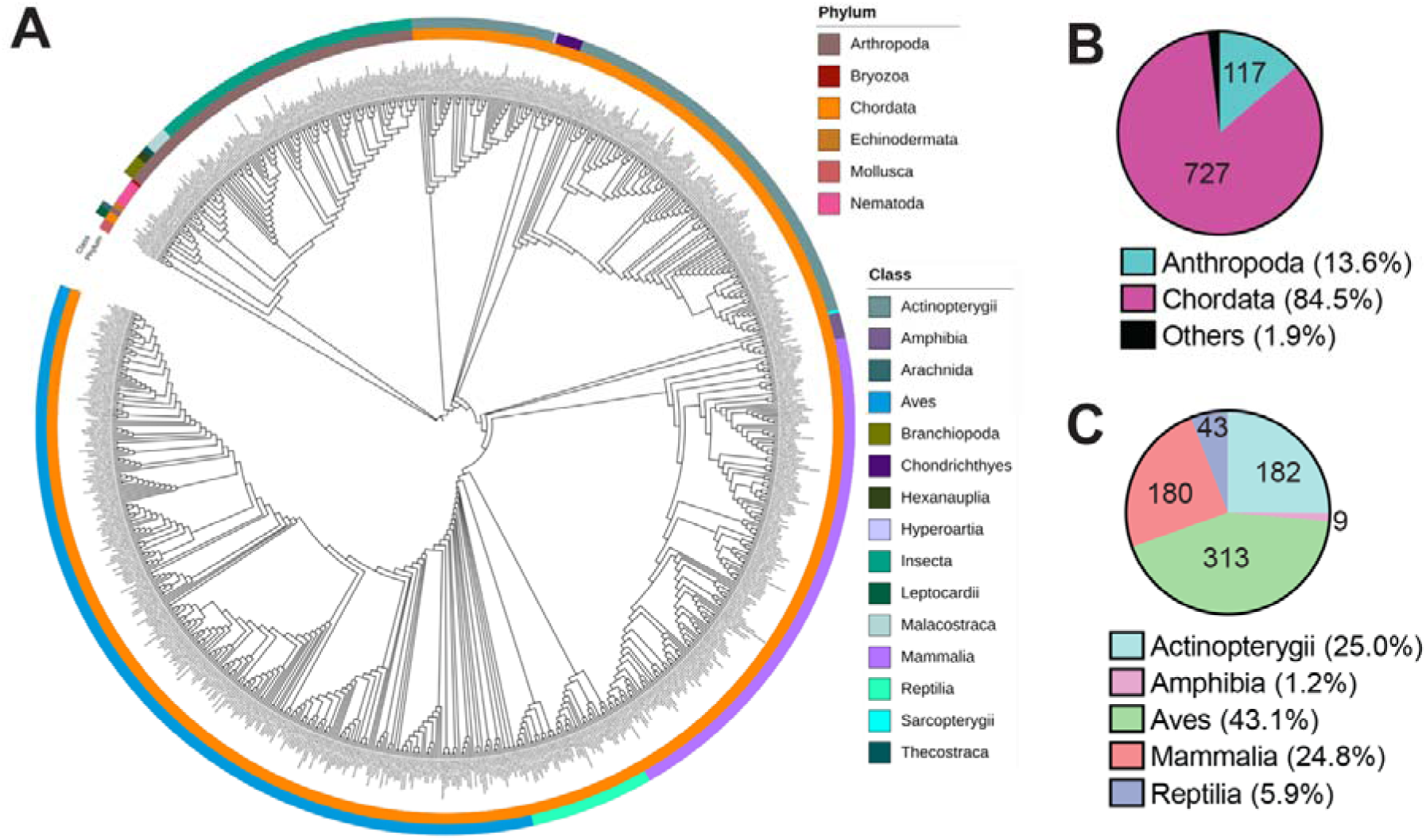
Phylogenetic analysis of ABHD12 sequences. (**A**) Phylogenetic tree representing the identified ABHD12 sequences from 860 organisms. The outer and inner colored circles represent the *Class* and *Phylum* respectively, to which a sequence belongs. (**B, C**) Pie-chart analysis representing the data from the phylogenetic tree from (**B**) different *Phyla*, and (**C**) various *Classes* within *phylum Chordata*. The number on the pie-chart represents number of ABHD12 sequences within that respective category. The pie-chart analysis shows that *phylum Chordata* contains most of the ABHD12 sequences, that has major distribution in class *Aves*, *Actinopterygii* and *Mammalia*.

### Sequence conservation within ABHD12 protein sequences

To determine the extent of conservation in ABHD12 sequences across various organisms, we performed a multiple sequence alignment analysis on the 860 identified ABHD12 orthologs. From this analysis, we found that 96 residues (out of the 398 residues from human ABHD12) were positionally conserved at a frequency of > 90% across all the 860 identified ABHD12 orthologs (**Table S2**), showing that the overall protein sequence conservation across all organisms is ∼ 24%. Next, as previously reported by us^20^, we formed five distinct groups of functionally conserved amino acid residues: e.g. acidic amino acids, basic amino acids, amino acids containing heteroatoms, aromatic amino acids, and hydrophobic amino acids (**Table S2**), and assessed the aligned sequences for functionally conserved residues. Based on this analysis, we found that another 69 residues (out of the 398 residues from human ABHD12) were functionally conserved across the identified ABHD12 orthologs (**Table S2**), suggesting that the overall conservation in ABHD12 sequences across various organisms was high (∼ 42%). Next, a closer inspection of identified ABHD12 sequences within a particular class showed that within a particular class (e.g. *Mammalia*), there was a very-high sequence conservation of ABHD12 (> 80%), and an ABHD12 sequence from that class might serve as a representative example for the entire class. Across different classes within a particular phylum (e.g. *Chordata*), the overall sequence conservation was found to be about 60% for the identified ABHD12 sequences in that phylum. Of note, our sequence conservation analysis shows that several missense mutations in ABHD12 that are associated with the human neurological disorder PHARC (i.e. P87T, T125M, R186P, T202I, T253R, H372Q, L385P)^39–41^, occur in residues that are highly conserved across ABHD12 sequences from all organisms (**Table S2**), and perhaps, highlight the importance of these residues in the biochemical activity of ABHD12.

### Structural assessment of the ABHD12 protein sequence

In the absence of an experimentally solved three-dimensional structure of ABHD12, to establish a structure-sequence relationship, we decided to use the structure of hABHD12 (UniProt ID: Q8N2K0) that was generated by the AlphaFold algorithm^29, 30^. To ensure that this predicted AlphaFold structure was reliable for our analysis, we overlaid this hABHD12 structure with an experimentally solved structure of human ABHD14B (PDB ID: 1imj)^37, 42, 43^ (**Figure S1**). From this analysis, we observed that despite being distantly related within the metabolic SH family^9^ (sequence identity <15%), the ABHD-folds of both these enzymes overlay well (RMSD ∼ 3.5 Å). Furthermore, we also found that the catalytic triad of the active sites (critical for catalysis) for both these enzymes had very similar orientations, poised for optimal catalysis (**Figure S1**). This congruence in the overall ABHD-fold and the active site orientations, despite the evolutionary distance, provided sufficient confidence that the predicted structure of hABHD12 would be useful for further structural analyses. Notably, the AlphaFold structure of hABHD12 also displayed an α-helical domain at the N-terminus of the protein sequence that presumably anchors this enzyme to the ER membrane, consistent with previous experimental observations^16, 19^. Next, we mapped all the conserved residues (including functionally conserved residues) on the hABHD12 AlphaFold structure (**Figure 4**). From this structural mapping, we found that most conserved residues clustered around the enzyme active site (catalytic triad) and the regions adjoining it, that comprise of a cleft and few flexible loops predicted to recognize and bind the substrate.

**Figure 4.**
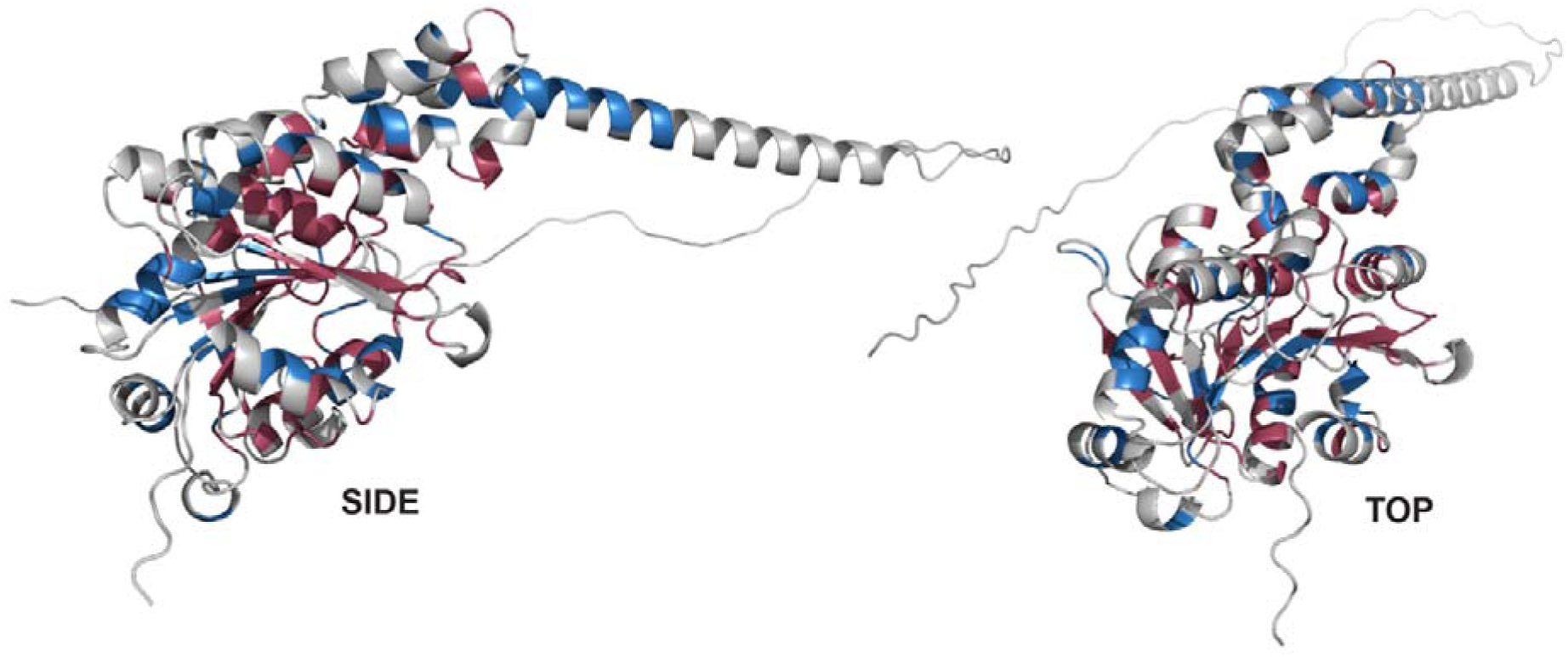
Mapping the conserved residues on the hABHD12 AlphaFold structure. Ribbon representation of the Alphafold2 predicted structure of hABHD12 (UniProt ID: Q8N2K0). Residues colored in red are fully conserved, while those shown in blue are functionally conserved based on the multiple sequence alignment analysis done on ABHD12 sequences from 860 different organisms.

To identify the putative lyso-PS binding pocket of hABHD12 and understand how the substrate accesses the active site, we used two computational tools (CAVER and MOLE 2.5) to map possible tunnels in the protein. Both the tools computed multiple tunnels in hABHD12, but only one surface channel/tunnel predicted by both the software included the crucial catalytic serine residue (Ser-246) needed for catalysis (**Figure 5A**). This predicted catalytic pocket of hABHD12 comprises of 24 residues of which 19 are conserved (12 fully conserved + 7 functionally conserved) (**Table S1**). Further, electrostatic calculations on the hABHD12 structure shows a positively charged region in the catalytic pocket, distal to the transmembrane helix (**Figure S2**). This region might facilitate the binding and orientation of the negatively charged lyso-PS head group within the active site. Interestingly, Ser-246 is located at the bottom of a deep negatively charged pocket adjacent to this transmembrane helix, and perhaps negative charges push the carbonyl oxygen of the ester of lyso-PS away from the catalytic site, facilitating the nucleophilic attack by Ser-246 (**Figure S2**). Finally, molecular docking of C18:0 lyso-PS on hABHD12 indicates that the substrate interacts closely with Asn-177, Glu-281 and His-285 (all fully conserved) and the side chains of some hydrophobic residues (most of which are conserved) in the 9 predicted ligand-binding conformations of C18:0 lyso-PS (**Figure 5B**). In addition to these, our modeling of C18:0 lyso-PS on hABHD12 also suggests that it makes polar contacts with charged residues located outside the catalytic pocket, namely His-185, Arg-186, His-245, His-372 and Lys-373 (all fully conserved) (**Figure 5B**).

**Figure 5.**
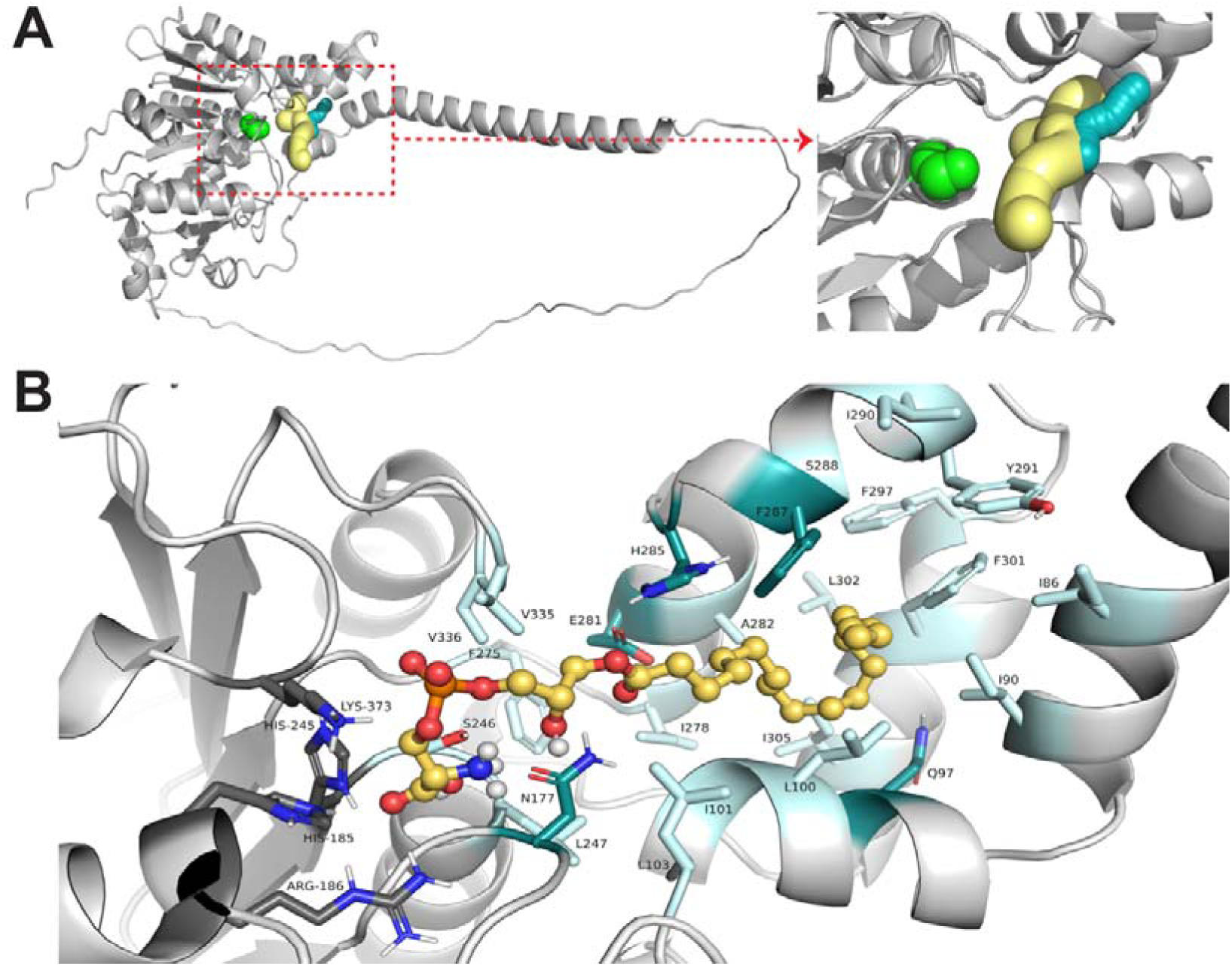
The lyso-PS binding pocket in hABHD12. (**A**) Identification of a lyso-PS binding pocket using two computational tools (CAVER and MOLE 2.5) (*left*), and a zoomed in version of this lyso-PS binding pocket (*right*), showing its proximity to the active site serine (Ser-246) residue (green spheres). (**B**) Mapping the putative lyso-PS interacting residues on hABHD12 with C18:0 lyso-PS, which is docked in the lyso-PS binding pocket. The residues colored dark green are predicted to directly interact with lyso-PS and are within the catalytic pocket. The residues in dark grey are also predicted to interact with lyso-PS but lie outside the catalytic pocket. Residues highlighted in light blue are part of the catalytic pocket.

Lastly, as part of this structural analysis, we also mapped the various missense PHARC mutants (P87T, T125M, R186P, T202I, T253R, H372Q, L385P)^39–41^ on the hABHD12 AlphaFold structure (**Figure S3**). Quite interestingly, we found that most of these missense mutations are on conserved residues that are part of important structural elements, such as α-helices or β-strands, or near the active site. Based on this structural analysis, we think that these missense PHARC mutations most probably affect local structures, which in turn are likely to affect the stability and overall enzymatic activity of hABHD12.

### Biochemical characterization of mABHD12 variants

Based on the structural mapping of conserved residues and the modeling of lyso-PS in the active site of hABHD12, we decided to biochemically validate the role of some residues in the enzymatic (lyso-PS lipase) activity. The hABHD12 and mABHD12 homologs are nearly identical in protein sequences (sequence identify 94%), and all residues discussed earlier are completely conserved between these two sequences at the very same position (**Figure S4**). Since we have previously established the overexpression and activity assays with recombinant mABHD12 from mammalian HEK293T cell membrane proteomes, coupled with the high sequence similarity between the two orthologs, we decided to perform all biochemical validations using mABHD12^16, 17^. From the bioinformatics and structural analysis, we shortlisted 22 conserved residues that were either part of the catalytic triad, proximal to the active site, or interactors of lyso-PS or residues having missense mutations in human PHARC subjects and mutated these to either alanine or the corresponding pathogenic PHARC variants for further biochemical studies (**Table 1**). All mutants were individually assayed using established gel-based ABPP assays and LC-MS based lyso-PS lipase assays to determine the nucleophilicity of the catalytic serine (Ser-246) and its ability to turn over the lyso-PS substrate respectively.

**Table 1:**
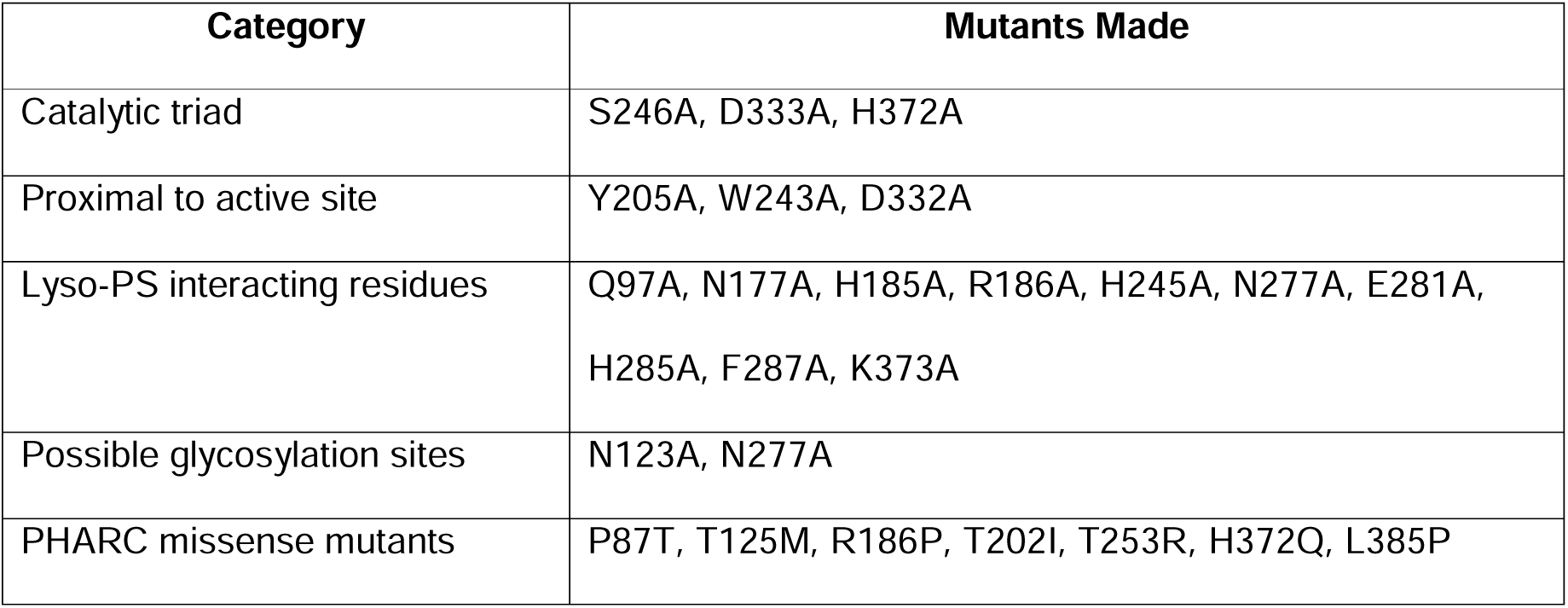
List of conserved residues selected for mutation and biochemical assays.

| Category | Mutants Made |
| --- | --- |
| Catalytic triad | S246A, D333A, H372A |
| Proximal to active site | Y205A, W243A, D332A |
| Lyso-PS interacting residues | Q97A, N177A, H185A, R186A, H245A, N277A, E281A, H285A, F287A, K373A |
| Possible glycosylation sites | N123A, N277A |
| PHARC missense mutants | P87T, T125M, R186P, T202I, T253R, H372Q, L385P |

First, we assayed the mutants of the residues from the catalytic triad (S246A, D333A and H372A) and those proximal to the active site (Y205A, W243A and D332A). Gel-based ABPP assays showed that mutations to the residues from the catalytic triad or those proximal to the active site resulted in substantial loss of the nucleophilicity of Ser-246 relative to the WT control (**Figure 6A**). Consistent with the gel-based ABPP assays, we found that except for D333A, mutations to all the aforementioned residues from the catalytic triad or those proximal to the active site resulted in almost complete loss of the lyso-PS lipase activity relative to the WT control (**Figure 6B**). Interestingly, we found that while the D333A mutant resulted in the significant loss of nucleophilicity of Ser-246 based on gel-based ABPP assays (**Figure 6A**), it retained ∼ 20% WT mABHD12 lyso-PS lipase activity **Figure 6B**), suggesting that the catalytic dyad of Ser-246 and His-372, were semi-competent in performing the enzymatic activity.

**Figure 6.**
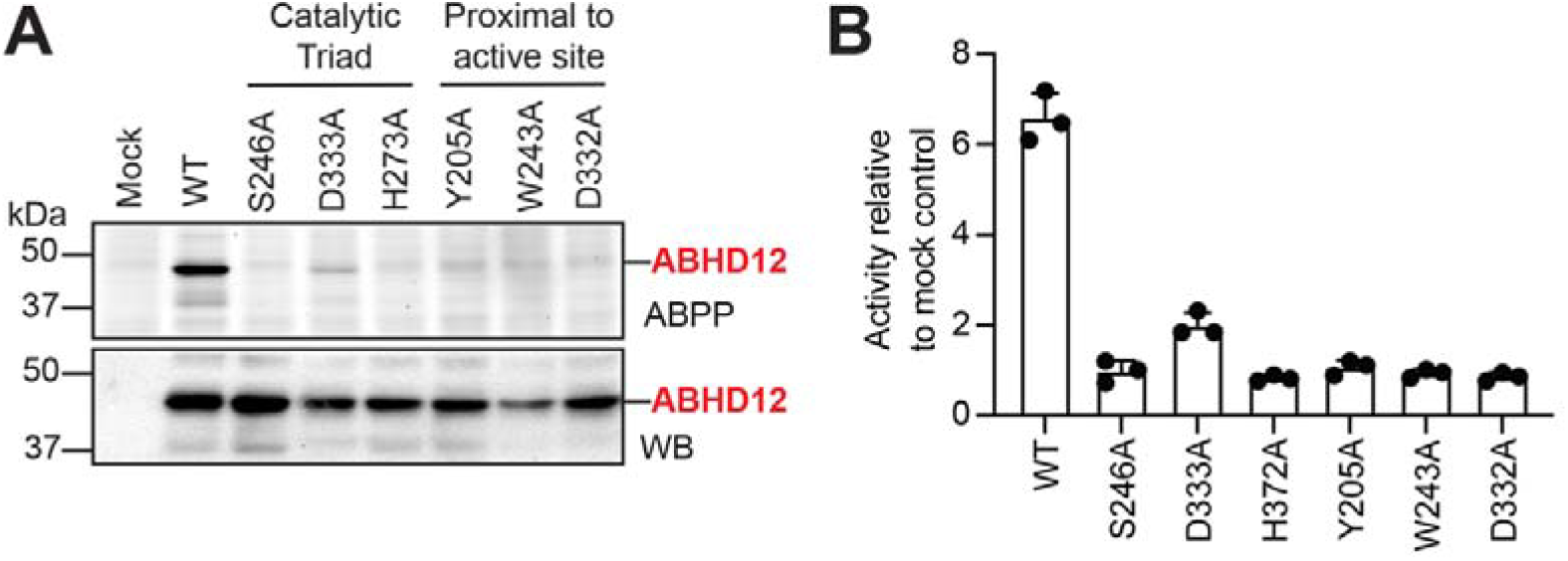
Biochemical characterization of mutants of the catalytic triad and residues proximal to the active site of mABHD12. (**A**) Membrane proteomic fractions of HEK293T cells transfected with mock, WT mABHD12, or various mutants of the catalytic triad (S246A, D333A, H372A) or conserved residues proximal to the active site (Y205A, W243A, D332A) were assessed by gel-based ABPP (top panel), and Western blot (WB) analysis using an anti-ABHD12 antibody (bottom panel). The band corresponding to ABHD12 (in red) represents the activity (top panel) and expression (bottom panel) of mABHD12 variants in membrane proteomic fractions of HEK293T cells. This experiment was done three times with reproducible results each time. (**B**) The lyso-PS lipase activity shown by membrane proteomic fractions of HEK293T cells transfected with mock, WT mABHD12, or various mutants of the catalytic triad (S246A, D333A, H372A) or conserved residues proximal to the active site (Y205A, W243A, D332A). The data was normalized to the activity of the mock control. The bars represent the mean ± standard deviation from three biological replicates per group.

Next, we assayed all the mutants of the residues predicted to interact with lyso-PS (based on the docking studies) (Q97A, N177A, H185A, R186A, H245A, N277A, E281A, H285A, F287A, K373A), and the possible glycosylation sites (N123A, N277A). We have previously shown that glycosylation is critical for ABHD12 activity^16^, and hence, we were interested in mapping the glycosylation site on ABHD12. Gel-based ABPP assays showed that while both mutants (N123A, N277A) showed diminished nucleophilicity of Ser-246, the N123A mutant, but not the N277A mutant, migrated significantly lower on the gel (based on the Western blot analysis) relative to the WT control, consistent with a non-glycosylated variant of mABHD12 (**Figure 7A**). Further, lyso-PS lipase activity assays showed that both mutants (N123A, N277A) had almost complete loss of this enzymatic activity relative to the WT control (**Figure 7B**), suggesting that Asn-123 is likely the site of glycosylation on ABHD12, while Asn-277 is an important lyso-PS interacting residue on ABHD12. Further, gel-based ABPP assays showed that amongst the various mutants for the lyso-PS interacting residues, Q97A, N177A, H185A, H285A, F287A, and K373A showed almost WT equivalent nucleophilicity of Ser-246 (**Figure 7A**). Quite surprisingly, the E281A mutant showed substantially more activity relative to the WT control in gel-based ABPP assays (**Figure 7A**). Amongst these lyso-PS interacting residue mutants, R186A and H245A (along with N277A) showed no discernable activity relative to the WT control (**Figure 7A**). Interestingly, while several mutants showed that the nucleophilicity of the active site Ser-246 was comparable to the WT control, lyso-PS lipase assays showed that only E281A mutant (∼ 90% of WT control), and to a lesser extent Q97 and F287A (∼ 50% of WT control), had this enzymatic activity (**Figure 7B**). Mutations to all other lyso-PS interacting residues resulted in an almost complete loss of lyso-PS lipase activity, suggesting their importance in binding lyso-PS (**Figure 7B**).

**Figure 7.**
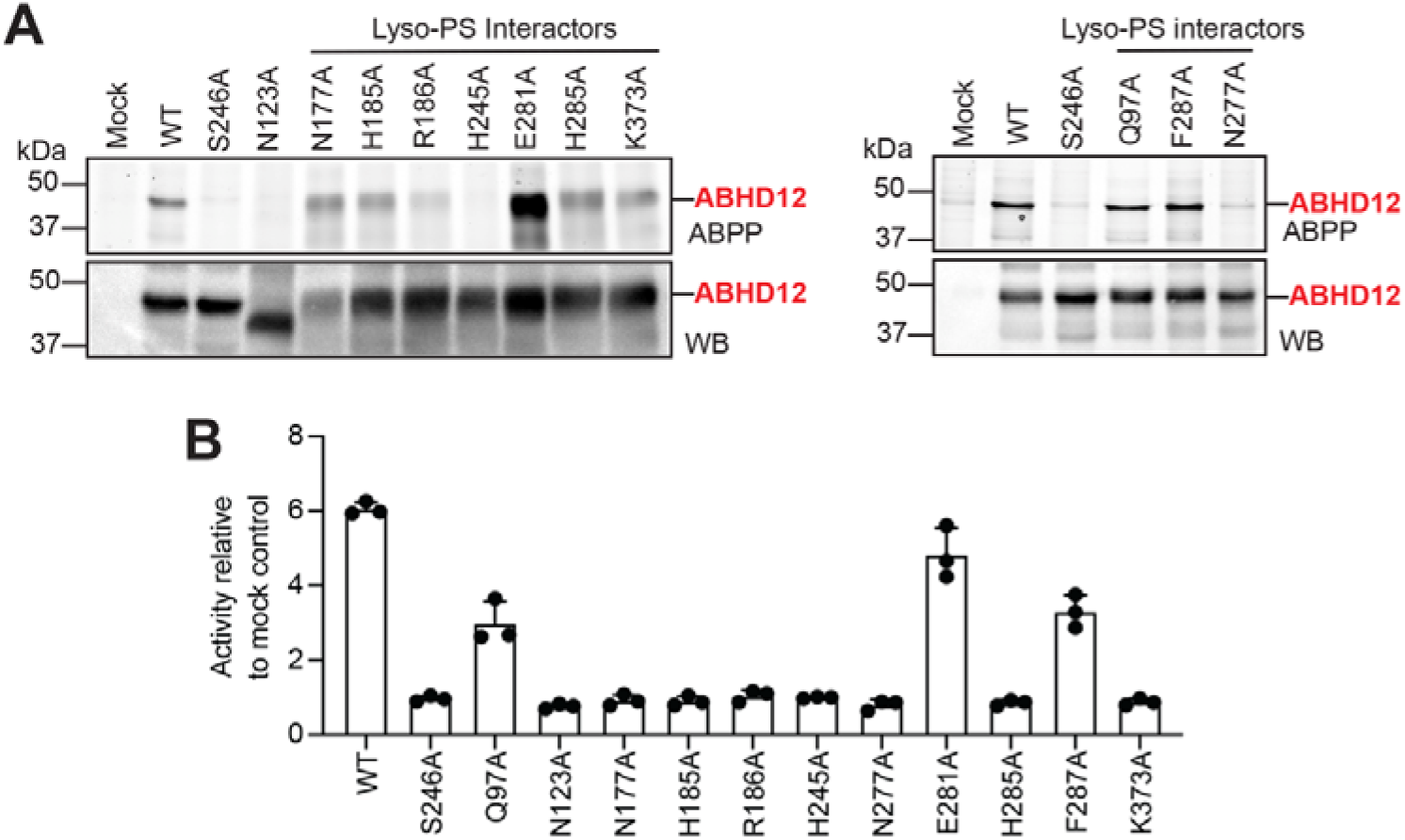
Biochemical characterization of mutants of residues predicted to bind lyso-PS or involved in glycosylation of mABHD12. (**A**) Membrane proteomic fractions of HEK293T cells transfected with mock, WT mABHD12, S246A mABHD12, or various mutants of residues predicted to bind/interact with lyso-PS (Q97A, N177A, H185A, R186A, H245A, N277A, E281A, H285A, F287A, K373A) or involved in glycosylation of mABHD12 (N123A, N277A) were assessed by gel-based ABPP (top panel), and Western blot (WB) analysis using an anti-ABHD12 antibody (bottom panel). The band corresponding to ABHD12 (in red) represents the activity (top panel) and expression (bottom panel) of mABHD12 variants in membrane proteomic fractions of HEK293T cells. This experiment was done three times with reproducible results each time. (**B**) The lyso-PS lipase activity shown by membrane proteomic fractions of HEK293T cells transfected with mock, WT mABHD12, S246A mABHD12, or various mutants of residues predicted to bind/interact with lyso-PS (Q97A, N177A, H185A, R186A, H245A, N277A, E281A, H285A, F287A, K373A) or involved in glycosylation of mABHD12 (N123A, N277A). The data was normalized to the activity of the mock control. The bars represent the mean ± standard deviation from three biological replicates per group. In both (**A**) and (**B**), the S246A mutant was used as an additional no activity controls in the assays.

Finally, we assayed all the missense mutants reported from human PHARC subjects (P87T, T125M, R186P, T202I, T253R, H372Q, L385P) using both the gel-based ABPP (**Figure 8A**) and lyso-PS lipase activity (**Figure 8B**) assays. Not surprisingly, we found that both the complementary assays showed that all the missense mutants reported from human PHARC subjects (P87T, T125M, R186P, T202I, T253R, H372Q, L385P) showed no discernable activity relative to the WT control. His-372 is part of the catalytic triad (**Figure 6**), and Arg-186 is an important residue involved in binding lyso-PS (**Figure 7**), and hence, together with our biochemical studies, missense mutations of these residues lacking enzymatic activity is expected. Based on the AlphaFold structure of hABHD12, we find that Pro-87, Thr-202, Thr-253 and Leu-385 are all highly conserved residues that are present on structurally conserved α-helices (**Figure S3**). We hypothesize that the missense mutations to these residues (P87T, T202I, T253R, L385P) most likely destabilize the local structure, and perhaps, results in the loss of enzymatic activity of ABHD12. Amongst the missense mutations, interestingly, we find by Western blot analysis that T125M migrates significantly lower on the gel (**Figure 8A**), and appears to exist predominantly in the non-glycosylated form of ABHD12. Given its proximity to Asn-123 (the putative glycosylation site on ABHD12), the T125M mutation likely disrupts the canonical glycosylation site (N-x-S/T) of ABHD12, and by doing so, hampers the enzymatic activity.

**Figure 8.**
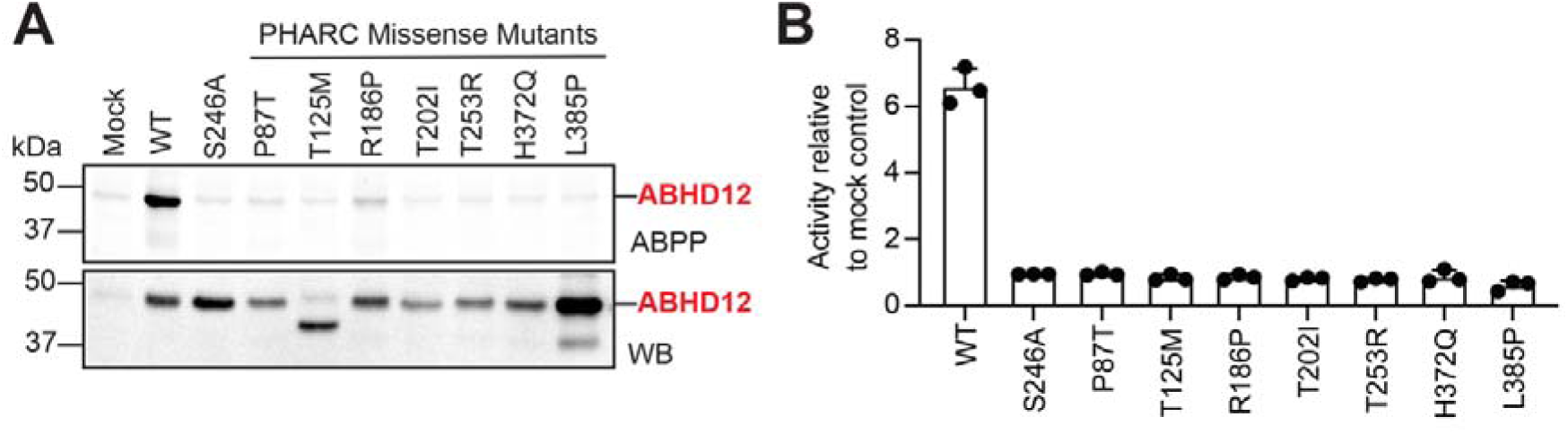
Biochemical characterization of missense mutations from human PHARC subjects. (**A**) Membrane proteomic fractions of HEK293T cells transfected with mock, WT mABHD12, S246A mABHD12, or various mutants of missense mutations from human PHARC subjects (P87T, T125M, R186P, T202I, T253R, H372Q, L385P) were assessed by gel-based ABPP (top panel), and Western blot (WB) analysis using an anti-ABHD12 antibody (bottom panel). The band corresponding to ABHD12 (in red) represents the activity (top panel) and expression (bottom panel) of mABHD12 variants in membrane proteomic fractions of HEK293T cells. This experiment was done three times with reproducible results each time. (**B**) The lyso-PS lipase activity shown by membrane proteomic fractions of HEK293T cells transfected with mock, WT mABHD12, S246A mABHD12, or various mutants of missense mutations from human PHARC subjects (P87T, T125M, R186P, T202I, T253R, H372Q, L385P). The data was normalized to the activity of the mock control. The bars represent the mean ± standard deviation from three biological replicates per group. In both (**A**) and (**B**), the S246A mutant was used as an additional no activity controls in the assays.

### Lyso-PS lipase activity of CG15111

To understand the physiological and functional role that ABHD12 activity plays in the context of PHARC, two animal models, namely mouse (*Mus musculus*)^11^ and zebrafish (*Danio rerio*)^41^, have been reported. Over the years, *Drosophila melanogaster* (common fruit fly) has also served as an excellent animal model for studying various human neurodegenerative diseases^44–48^, given the ease of genetically manipulating them^49–52^. Our bioinformatics and phylogenetic analysis identified an uncharacterized lipase CG15111 from the metabolic SH family in *D. melanogaster* as an ortholog of hABHD12 (sequence identity 37%)^36^ (**Figure S4**). To test if CG15111 possesses any lyso-PS lipase activity similar to that of mammalian ABHD12, we cloned and expressed wild-type (WT) CG15111 in insect (S2) cells using established protocols. Bioinformatics analysis suggested that Ser-241 of CG15111 is the catalytic nucleophile^36^, and hence we made the S241A CG15111 mutant, and also expressed this in insect cells, as a negative control for the biochemical assays with CG15111.

By Western blot analysis, we found that relative to the “mock” control, both WT CG15111 and S241A CG15111 showed comparable overexpression in the membrane proteomic lysates of insect cells (**Figure 9A**). Next, using gel-based ABPP assays, we show that WT CG15111, but not the S241A CG15111 mutant, is active in the membrane proteomic lysates of insect cells (**Figure 9A**). Furthermore, to ascertain if CG15111 can hydrolyze lyso-PS, we assayed all the aforementioned membrane proteomic lysates of insect cells, namely mock, WT CG15111, and S241A CG15111 for lyso-PS lipase activity. From this experiment we found that relative to “mock” control lysates, WT CG15111 had robust lyso-PS lipase activity, while the S241A CG15111 had negligible enzymatic activity for the same reaction (**Figure 9B**). Taken together, these results confirm the bioinformatics prediction that CG15111 is indeed an ortholog of mammalian ABHD12 that possesses robust lyso-PS lipase activity, and Ser-241 of CG15111 is the conserved nucleophilic serine residue critical for catalysis.

**Figure 9.**
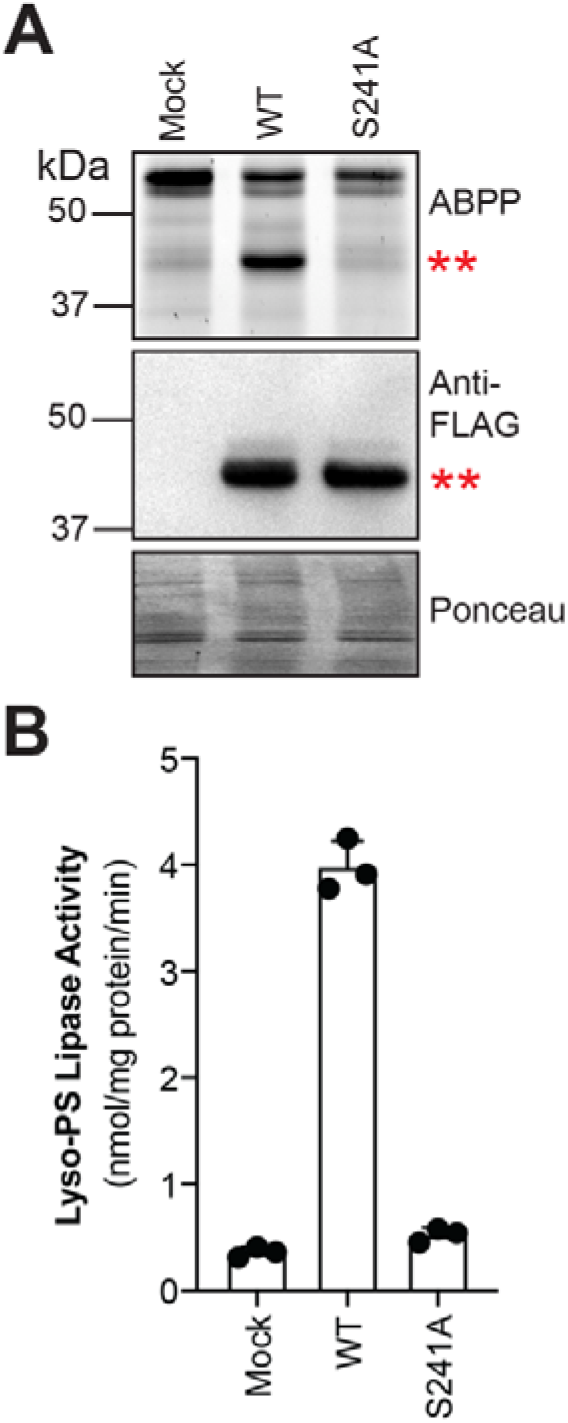
Enzymatic activity of CG15111 expressed in S2 cells. (**A**) Membrane proteomic fractions of S2 cells transfected with mock, WT CG15111, or S241A CG15111 were assessed by gel-based ABPP (top panel), Western blot analysis using an anti-FLAG antibody (middle panel), and Ponceau S staining (bottom panel). The double red asterisk represents the activity (top panel) and expression (middle panel) of CG15111 in membrane proteomic fractions of S2 cells. This experiment was done three times with reproducible results each time. (**B**) The lyso-PS lipase activity shown by membrane proteomic fractions of S2 cells transfected with mock, WT CG15111, or S241A CG15111. The data shows that WT CG15111, but not S241A CG15111, has robust lyso-PS lipase activity. The bars represents the mean ± standard deviation from three biological replicates per group.

## DISCUSSION

Advances in DNA sequencing technologies have greatly facilitated our understanding of the genetic basis of hereditary human disorders^53^. Of note, to date, genome mapping and sequencing efforts from human subjects suffering from a variety of clinical symptoms have led to the identification of >4500 inherited human diseases (catalogued in the OMIM database^3–5^), with newer pathogenic mutations still being discovered at a rapid rate. As newer disease-causing mutations continue to be mapped in humans, it is becoming apparent that many of the affected genes code for proteins that are uncharacterized and/or have poorly understood biochemical function^54^. Hence, mechanistically characterizing the biochemical functions of such proteins is going to be critical to achieve a deeper mechanistic understanding of human genetic disorders and for identifying potential strategies for treating them. The neurological disorder PHARC serves as a nice example of this premise in the context of human hereditary diseases^6, 7^. In 2010, genome sequencing efforts led to the identification of PHARC as an autosomal recessive genetic disorder in humans caused by deleterious mutations in the *abhd12* gene^1, 2^. Since this discovery, over the past decade, >30 distinct homozygous or compound heterozygous mutations (deletion-insertion, nonsense, frameshift, splice site, and missense type mutations) in the *abhd12* gene have now been identified, all leading to the symptoms associated with PHARC in humans^6, 7^. The *abhd12* gene encodes an integral membrane associated lipase ABHD12 from the metabolic SH family, that physiologically functions as the major lyso-PS lipase in the mammalian central nervous and immune systems (**Figure 1**, **Figure 2**)^11, 12, 14–17^. While mutations (e.g. deletion-insertion, nonsense, frameshift, and splice site) leading to the truncated full-length ABHD12 protein in the context of PHARC have been easy to reconcile in terms of the enzymatic activity of this lipase, the various missense mutations associated with PHARC remain difficult to biochemically explain in the absence of any detailed protein sequence-enzyme activity relationship studies for ABHD12.

To overcome this knowledge gap and develop a protein sequence-enzyme activity relationship profile for ABHD12, we first performed a thorough bioinformatics analysis and collated all putative ABHD12 sequences from all sequenced organisms across the evolutionary time scale (**Figure 3, Table S2**). Next, complementarily using multiple sequence alignment analysis and the AlphaFold structure of hABHd12, we generated a map of the structurally conserved residues across all ABHD12 sequences, and found that a significant portion of the overall ABHD12 structure is conserved across all organisms (**Figure 4, Table S2**). Further, using this information along with molecular docking of C18:0 lyso-PS into the ABHD12, we identify a hydrophobic cavity near the active site that is capable of binding lyso-PS, and identify putative lyso-PS interacting residues from this analysis (**Figure 5**). Following up on all the *in-silico* analysis, using gel-based ABPP and LC-MS based lyso-PS lipase activity assays, we biochemically assay several ABHD12 mutants. These ABHD12 mutants comprise of residues that are part of the catalytic triad or proximal to the active site (**Figure 6**), or residues that interact with and enable binding of lyso-PS (**Figure 7**) or missense mutants that are found in human PHARC subjects (**Figure 8**). Together, these studies establish the first exhaustive sequence-enzyme activity relationship for ABHD12, and explain why missense mutations found in human PHARC subjects lack lyso-PS lipase activity. Finally, our evolutionary analysis identifies a fly ortholog of hABHD12, namely CG15111. We recombinantly express this fly ortholog of hABHD12 and show that CG15111 has robust lyso-PS lipase activity (**Figure 9**).

Moving forward, firstly, the bioinformatics and the sequence-enzyme activity relationship data generated in this study might prove to be a valuable predictive tool in understanding why emerging missense mutations in ABHD12 associated with human PHARC subjects result in diminished lyso-PS lipase activity. Secondly, we have previously shown that ABHD12 needs to be glycosylated for optimal activity^16^, and prefers VLC lyso-PSs as substrates^17^. While there is enough biochemical evidence in support of the need for glycosylation and the substrate preference of ABHD12, the AlphaFold predicted structure of hABHD12 and molecular docking studies cannot fully explain both of these. Hence, solving the structure of ABHD12 experimentally might be needed to corroborate these biochemical findings and understand in more detail the substrate preference and the structural effect that glycosylation has on the enzymatic activity of ABHD12. Lastly, our identification of CG15111, the fly ortholog of hABHD12, opens several new research avenues to biochemically explore how dysregulated ABHD12 activity might contribute to PHARC. For example, given the lifespan of flies (relative to mice models), understanding the age-dependent progression and neurobehavioral defects associated with PHARC can be studied in a quicker time scale, particularly in the context of the missense mutations found in human PHARC subjects. Towards this, it would also be interesting to see if genetic tools (such as CRISPR/Cas9) can be used in flies to first model the missense mutations found in human PHARC subjects, and then, if these mutations can also be reversed (or rescued) using the same genetic approaches. Such studies from fly models in the future may hold promise in personalized gene therapy for treating PHARC and perhaps, even other similar hereditary human diseases.

## DATA AVAILABILITY

All the data that supports the findings reported in this study are available in the paper and its associated supporting information or are available with the corresponding author upon reasonable request.

## ACCESSION CODES

The UniProt ID of human ABHD12 is Q8N2K0, mouse ABHD12 is Q99LR1 and *Drosophila melanogaster* ABHD12 (CG15111) is A1ZBH2. The NCBI accession IDs of all ABHD12 homologs identified from the bioinformatics analysis can be found in **Table S2**.

## SUPPORTING INFORMATION

The Supporting Information associated with this paper is available free of cost, and contains **Figures S1 – S4**, and **Table S1 – S2**.

## AUTHOR CONTRIBUTIONS

A.C. performed all the biochemical assays with assistance from A.D., A.D. performed all the bioinformatics and structural analysis with M.S.M., K.K., and R.C.S. cloned and expressed CG15111 used in this study under the supervision of G.S.R., S.S.K. supervised the project and acquired funding for the project. A.C., A.D., and S.S.K. wrote the paper with inputs from all authors.

## ACKNOWLEDGEMENTS

Members of the S.S.K. and G.S.R. lab are thanked for valuable comments and discussions on this study. The authors thank Golding Rodrigues for advice on the bioinformatics analysis with ABHD12, and Saddam Shekh for technical assistance and maintenance of the biological mass spectrometry facility at IISER Pune. Dr. Kavita Sharma is thanked for synthesizing the FP-probes used in this study.

## REFERENCES

[1] Fiskerstrand, T., H’Mida-Ben Brahim, D., Johansson, S., M’Zahem, A., Haukanes, B. I., Drouot, N., Zimmermann, J., Cole, A. J., Vedeler, C., Bredrup, C., Assoum, M., Tazir, M., Klockgether, T., Hamri, A., Steen, V. M., Boman, H., Bindoff, L. A., Koenig, M., and Knappskog, P. M. (2010) Mutations in ABHD12 cause the neurodegenerative disease PHARC: An inborn error of endocannabinoid metabolism, Am J Hum Genet. 87, 410–417.

[2] Fiskerstrand, T., Knappskog, P., Majewski, J., Wanders, R. J., Boman, H., and Bindoff, L. A. (2009) A novel Refsum-like disorder that maps to chromosome 20, Neurology 72, 20–27.

[3] Amberger, J. S., Bocchini, C. A., Schiettecatte, F., Scott, A. F., and Hamosh, A. (2015) OMIM.org: Online Mendelian Inheritance in Man (OMIM(R)), an online catalog of human genes and genetic disorders, Nucleic Acids Res 43, D789–798.

[4] Hamosh, A., Scott, A. F., Amberger, J., Valle, D., and McKusick, V. A. (2000) Online Mendelian Inheritance in Man (OMIM), Hum Mutat 15, 57–61.

[5] Schorderet, D. F. (1991) Using OMIM (On-line Mendelian Inheritance in Man) as an expert system in medical genetics, American journal of medical genetics 39, 278–284.

[6] Daneshi, A., Garshasbi, M., Farhadi, M., Falavarjani, K. G., Vafaee-Shahi, M., Almadani, N., Zabihi, M., Ghalavand, M. A., and Falah, M. (2023) Genetic insights into PHARC syndrome: identification of a novel frameshift mutation in ABHD12, Bmc Med Genomics 16.

[7] Harutyunyan, L., Callaerts, P., and Vermeer, S. (2024) PHARC syndrome: an overview, Orphanet journal of rare diseases 19, 416.

[8] Eisenberger, T., Slim, R., Mansour, A., Nauck, M., Nurnberg, G., Nurnberg, P., Decker, C., Dafinger, C., Ebermann, I., Bergmann, C., and Bolz, H. J. (2012) Targeted next-generation sequencing identifies a homozygous nonsense mutation in ABHD12, the gene underlying PHARC, in a family clinically diagnosed with Usher syndrome type 3, Orphanet journal of rare diseases 7, 59.

[9] Long, J. Z., and Cravatt, B. F. (2011) The metabolic serine hydrolases and their functions in mammalian physiology and disease, Chem Rev 111, 6022–6063.

[10] Blankman, J. L., Simon, G. M., and Cravatt, B. F. (2007) A comprehensive profile of brain enzymes that hydrolyze the endocannabinoid 2-arachidonoylglycerol, Chem Biol 14, 1347–1356.

[11] Blankman, J. L., Long, J. Z., Trauger, S. A., Siuzdak, G., and Cravatt, B. F. (2013) ABHD12 controls brain lysophosphatidylserine pathways that are deregulated in a murine model of the neurodegenerative disease PHARC, Proc Natl Acad Sci U S A 110, 1500–1505.

[12] Chakraborty, A., and Kamat, S. S. (2024) Lysophosphatidylserine: A Signaling Lipid with Implications in Human Diseases, Chem Rev 124, 5470–5504.

[13] Shanbhag, K., Sharma, K., and Kamat, S. S. (2023) Photoreactive bioorthogonal lipid probes and their applications in mammalian biology, RSC Chem Biol 4, 37–46.

[14] Kamat, S. S., Camara, K., Parsons, W. H., Chen, D. H., Dix, M. M., Bird, T. D., Howell, A. R., and Cravatt, B. F. (2015) Immunomodulatory lysophosphatidylserines are regulated by ABHD16A and ABHD12 interplay, Nat Chem Biol 11, 164–171.

[15] Kelkar, D. S., Ravikumar, G., Mehendale, N., Singh, S., Joshi, A., Sharma, A. K., Mhetre, A., Rajendran, A., Chakrapani, H., and Kamat, S. S. (2019) A chemical-genetic screen identifies ABHD12 as an oxidized-phosphatidylserine lipase, Nat Chem Biol 15, 169–178.

[16] Joshi, A., Shaikh, M., Singh, S., Rajendran, A., Mhetre, A., and Kamat, S. S. (2018) Biochemical characterization of the PHARC-associated serine hydrolase ABHD12 reveals its preference for very-long-chain lipids, Journal of Biological Chemistry 293, 16953–16963.

[17] Khandelwal, N., Shaikh, M., Mhetre, A., Singh, S., Sajeevan, T., Joshi, A., Balaji, K. N., Chakrapani, H., and Kamat, S. S. (2021) Fatty acid chain length drives lysophosphatidylserine-dependent immunological outputs, Cell Chem Biol 28, 1169–1179.

[18] Singh, S., and Kamat, S. S. (2021) The loss of enzymatic activity of the PHARC-associated lipase ABHD12 results in increased phagocytosis that causes neuroinflammation, Eur J Neurosci 54, 7442–7457.

[19] Singh, S., Joshi, A., and Kamat, S. S. (2020) Mapping the Neuroanatomy of ABHD16A, ABHD12, and Lysophosphatidylserines Provides New Insights into the Pathophysiology of the Human Neurological Disorder PHARC, Biochemistry 59, 2299–2311.

[20] Vaidya, K., Rodrigues, G., Gupta, S., Devarajan, A., Yeolekar, M., Madhusudhan, M. S., and Kamat, S. S. (2023) Identification of sequence determinants for the ABHD14 enzymes, Proteins.

[21] Altschul, S. F., Madden, T. L., Schaffer, A. A., Zhang, J., Zhang, Z., Miller, W., and Lipman, D. J. (1997) Gapped BLAST and PSI-BLAST: a new generation of protein database search programs, Nucleic Acids Res 25, 3389–3402.

[22] Moller, S., Croning, M. D., and Apweiler, R. (2001) Evaluation of methods for the prediction of membrane spanning regions, Bioinformatics 17, 646–653.

[23] Dobson, L., Remenyi, I., and Tusnady, G. E. (2015) CCTOP: a Consensus Constrained TOPology prediction web server, Nucleic Acids Res 43, W408–412.

[24] Needleman, S. B., and Wunsch, C. D. (1970) A general method applicable to the search for similarities in the amino acid sequence of two proteins, J Mol Biol 48, 443–453.

[25] Sievers, F., Wilm, A., Dineen, D., Gibson, T. J., Karplus, K., Li, W., Lopez, R., McWilliam, H., Remmert, M., Soding, J., Thompson, J. D., and Higgins, D. G. (2011) Fast, scalable generation of high-quality protein multiple sequence alignments using Clustal Omega, Mol Syst Biol 7, 539.

[26] Tamura, K., Stecher, G., and Kumar, S. (2021) MEGA11: Molecular Evolutionary Genetics Analysis Version 11, Mol Biol Evol 38, 3022–3027.

[27] Jones, D. T., Taylor, W. R., and Thornton, J. M. (1992) The rapid generation of mutation data matrices from protein sequences, Comput Appl Biosci 8, 275–282.

[28] Letunic, I., and Bork, P. (2007) Interactive Tree Of Life (iTOL): an online tool for phylogenetic tree display and annotation, Bioinformatics 23, 127–128.

[29] Varadi, M., Anyango, S., Deshpande, M., Nair, S., Natassia, C., Yordanova, G., Yuan, D., Stroe, O., Wood, G., Laydon, A., Zidek, A., Green, T., Tunyasuvunakool, K., Petersen, S., Jumper, J., Clancy, E., Green, R., Vora, A., Lutfi, M., Figurnov, M., Cowie, A., Hobbs, N., Kohli, P., Kleywegt, G., Birney, E., Hassabis, D., and Velankar, S. (2022) AlphaFold Protein Structure Database: massively expanding the structural coverage of protein-sequence space with high-accuracy models, Nucleic Acids Res 50, D439–D444.

[30] Jumper, J., Evans, R., Pritzel, A., Green, T., Figurnov, M., Ronneberger, O., Tunyasuvunakool, K., Bates, R., Zidek, A., Potapenko, A., Bridgland, A., Meyer, C., Kohl, S. A. A., Ballard, A. J., Cowie, A., Romera-Paredes, B., Nikolov, S., Jain, R., Adler, J., Back, T., Petersen, S., Reiman, D., Clancy, E., Zielinski, M., Steinegger, M., Pacholska, M., Berghammer, T., Bodenstein, S., Silver, D., Vinyals, O., Senior, A. W., Kavukcuoglu, K., Kohli, P., and Hassabis, D. (2021) Highly accurate protein structure prediction with AlphaFold, Nature 596, 583–589.

[31] Stourac, J., Vavra, O., Kokkonen, P., Filipovic, J., Pinto, G., Brezovsky, J., Damborsky, J., and Bednar, D. (2019) Caver Web 1.0: identification of tunnels and channels in proteins and analysis of ligand transport, Nucleic Acids Research 47, W414–W422.

[32] Pravda, L., Sehnal, D., Tousek, D., Navratilova, V., Bazgier, V., Berka, K., Varekova, R. S., Koca, J., and Otyepka, M. (2018) MOLEonline: a web-based tool for analyzing channels, tunnels and pores (2018 update), Nucleic Acids Research 46, W368–W373.

[33] O’Boyle, N. M., Banck, M., James, C. A., Morley, C., Vandermeersch, T., and Hutchison, G. R. (2011) Open Babel: An open chemical toolbox, J Cheminform 3, 33.

[34] Bhaskar, V., Valentine, S. A., and Courey, A. J. (2000) A functional interaction between dorsal and components of the Smt3 conjugation machinery, J Biol Chem 275, 4033–4040.

[35] Kumar, K., Pazare, M., Ratnaparkhi, G. S., and Kamat, S. S. (2024) CG17192 is a Phospholipase That Regulates Signaling Lipids in the Gut upon Infection, Biochemistry.

[36] Kumar, K., Mhetre, A., Ratnaparkhi, G. S., and Kamat, S. S. (2021) A Superfamily-wide Activity Atlas of Serine Hydrolases in Drosophila melanogaster, Biochemistry 60, 1312–1324.

[37] Rajendran, A., Vaidya, K., Mendoza, J., Bridwell-Rabb, J., and Kamat, S. S. (2020) Functional Annotation of ABHD14B, an Orphan Serine Hydrolase Enzyme, Biochemistry 59, 183–196.

[38] Lord, C. C., Thomas, G., and Brown, J. M. (2013) Mammalian alpha beta hydrolase domain (ABHD) proteins: Lipid metabolizing enzymes at the interface of cell signaling and energy metabolism, Biochim Biophys Acta 1831, 792–802.

[39] Igelman, A. D., Ku, C., da Palma, M. M., Georgiou, M., Schiff, E. R., Lam, B. L., Sankila, E. M., Ahn, J., Pyers, L., Vincent, A., Sallum, J. M. F., Zein, W. M., Oh, J. K., Maldonado, R. S., Ryu, J., Tsang, S. H., Gorin, M. B., Webster, A. R., Michaelides, M., Yang, P., and Pennesi, M. E. (2021) Expanding the clinical phenotype in patients with disease causing variants associated with atypical Usher syndrome, Ophthalmic Genet 42, 664–673.

[40] Nishiguchi, K. M., Avila-Fernandez, A., van Huet, R. A., Corton, M., Perez-Carro, R., Martin-Garrido, E., Lopez-Molina, M. I., Blanco-Kelly, F., Hoefsloot, L. H., van Zelst-Stams, W. A., Garcia-Ruiz, P. J., Del Val, J., Di Gioia, S. A., Klevering, B. J., van de Warrenburg, B. P., Vazquez, C., Cremers, F. P., Garcia-Sandoval, B., Hoyng, C. B., Collin, R. W., Rivolta, C., and Ayuso, C. (2014) Exome sequencing extends the phenotypic spectrum for ABHD12 mutations: from syndromic to nonsyndromic retinal degeneration, Ophthalmology 121, 1620–1627.

[41] Tingaud-Sequeira, A., Raldua, D., Lavie, J., Mathieu, G., Bordier, M., Knoll-Gellida, A., Rambeau, P., Coupry, I., Andre, M., Malm, E., Moller, C., Andreasson, S., Rendtorff, N. D., Tranebjaerg, L., Koenig, M., Lacombe, D., Goizet, C., and Babin, P. J. (2017) Functional validation of ABHD12 mutations in the neurodegenerative disease PHARC, Neurobiology of disease 98, 36–51.

[42] Padmanabhan, B., Kuzuhara, T., Adachi, N., and Horikoshi, M. (2004) The crystal structure of CCG1/TAF(II)250-interacting factor B (CIB), J Biol Chem. 279, 9615–9624.

[43] Rajendran, A., Soory, A., Khandelwal, N., Ratnaparkhi, G., and Kamat, S. S. (2022) A multi-omics analysis reveals that the lysine deacetylase ABHD14B influences glucose metabolism in mammals, J Biol Chem 298, 102128.

[44] Chan, H. Y., and Bonini, N. M. (2000) Drosophila models of human neurodegenerative disease, Cell Death Differ 7, 1075–1080.

[45] McGurk, L., Berson, A., and Bonini, N. M. (2015) Drosophila as an In Vivo Model for Human Neurodegenerative Disease, Genetics 201, 377–402.

[46] Prüssing, K., Voigt, A., and Schulz, J. B. (2013) Drosophila melanogaster as a model organism for Alzheimer’s disease, Molecular neurodegeneration 8, 35.

[47] Marsh, J. L., and Thompson, L. M. (2006) Drosophila in the study of neurodegenerative disease, Neuron 52, 169–178.

[48] Muqit, M. M., and Feany, M. B. (2002) Modelling neurodegenerative diseases in Drosophila: a fruitful approach?, Nature reviews. Neuroscience 3, 237–243.

[49] Hales, K. G., Korey, C. A., Larracuente, A. M., and Roberts, D. M. (2015) Genetics on the Fly: A Primer on the Drosophila Model System, Genetics 201, 815–842.

[50] Rajan, A., and Perrimon, N. (2013) Of flies and men: insights on organismal metabolism from fruit flies, Bmc Biol 11 (38), 1–6.

[51] Bier, E. (2005) Drosophila, the golden bug, emerges as a tool for human genetics, Nat Rev Genet 6, 9–23.

[52] Bilder, D., and Irvine, K. D. (2017) Taking Stock of the Drosophila Research Ecosystem, Genetics 206, 1227–1236.

[53] Rabbani, B., Mahdieh, N., Hosomichi, K., Nakaoka, H., and Inoue, I. (2012) Next-generation sequencing: impact of exome sequencing in characterizing Mendelian disorders, J Hum Genet 57, 621–632.

[54] Galperin, M. Y., and Koonin, E. V. (2004) ’Conserved hypothetical’ proteins: prioritization of targets for experimental study, Nucleic Acids Res 32, 5452–5463.

