## Supplementary Figures for "Bioinformatics analysis identifies sequence determinants of enzymatic activity for the PHARC associated lipase ABHD12"

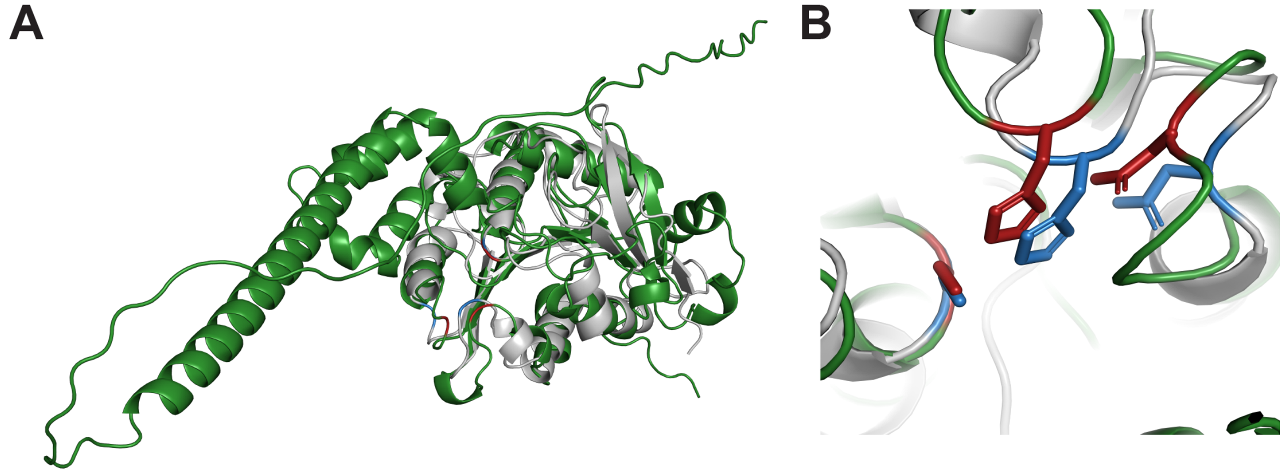


**Figure S1.** Structural alignment of hABHD12 and human ABHD14B. (**A**) Overlay of the experimentally solved structure of human ABHD14B (grey) (PDB: 1imj) and the AlphaFold predicted structure of hABHD12 (green) (UniProt ID: Q8N2K0), showing a good overall alignment of both structures, especially the ABHD-portion of the structure. (**B**) Overlay of the catalytic triad of human ABHD14B (blue) and hABHD12 (red), showing a near perfect orientation of the catalytic triad of the AlphaFold predicted structure of hABHD12.

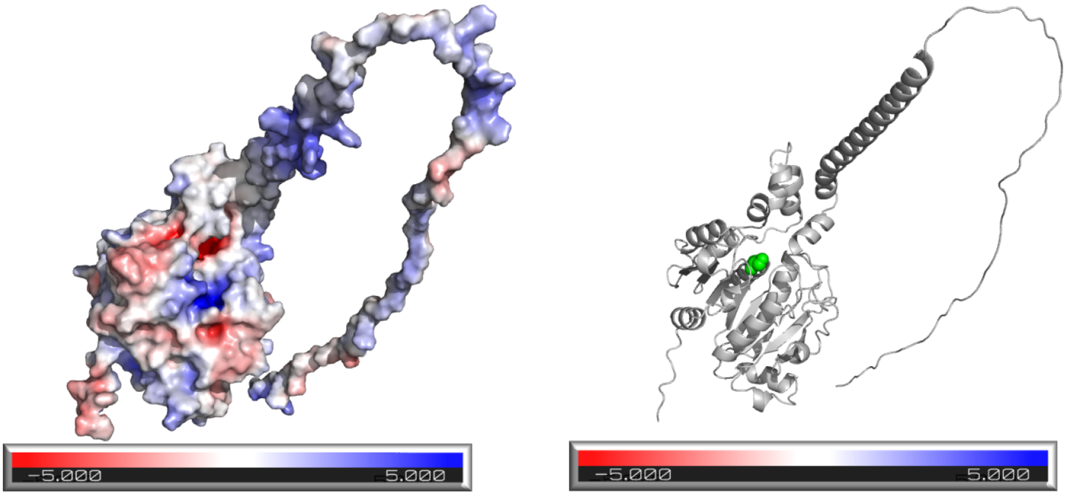


**Figure S2.** Electrostatic map of the predicted hABHD12 cavities. *Left*, the predicted electrostatic map of the various cavities on the AlphaFold predicted structure of hABHD12 identified using various software. *Right*, the same orientation of the structure of hABHD12 showing the position of the catalytic serine residue (green balls), to understand the electrostatic map in the context of the enzyme active site.

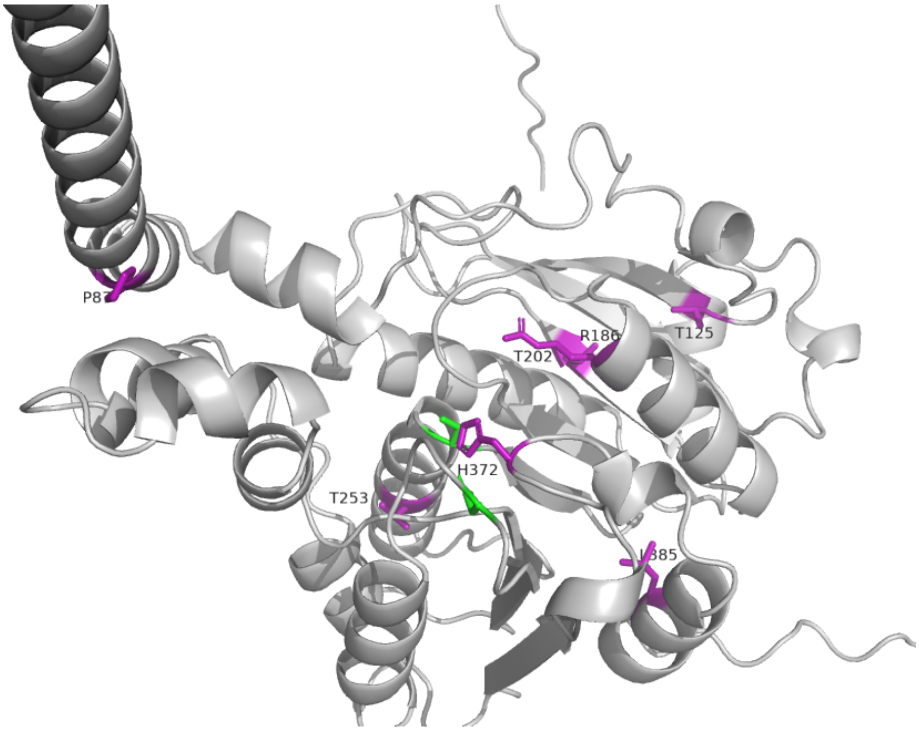


**Figure S3.** Mapping the locations of the identified missense PHARC mutations on the AlphaFold predicted structure of hABHD12 (green) (UniProt ID: Q8N2K0)

```
CG15111      -----------------------------------------------------------M      1
mABHD12      MRKRTEPVTLEHERCAASGSSSSGSAAAALDADCSLKQNLRLAGKGTAEPHSASDAGMKR      60
hABHD12      MRKRTEPVALEHERCAAAGSSSSGSAAAALDADCRLKQNLRLTGPAAAEPRCAADAGMKR      60


CG15111      LFTRRRLLKRVGLRCLQACLLIFFLIFVVLPLIFRYSVTFQRGILFLTFIKYPKGLDLTK      61
mABHD12      ALGRR-KSLWFRLRKILLCVLGF---YIAIPFLVKLCPGIQAKLIFLNFVRVPYFIDLKK      116
hABHD12      ALGRR-KGVWLRLRKILFCVLGL---YIAIPFLIKLCPGIQAKLIFLNFVRVPYFIDLKK      116
              : **     . ** :  *:* :   ::.:*::.: .  :*  ::**.*:: *  :**.*

CG15111      PESVGLYATRNFYITVKDHDQDEDGVRVGVWHVLPSNAVRRFKRELRVEEEVAQDPDQQL      121
mABHD12      PQDQGLNHTCNYYLQ------PEDDVTIGVWHTIPSVWWKNAQ-----------------      153
hABHD12      PQDQGLNHTCNYYLQ------PEEDVTIGVWHTVPAVWWKNAQ-----------------      153
             *:. **  * *:*:        *:.* :****.:*:   :. :

CG15111      DPAPGNERELKELSPAIRSEFPVVLPENEQLFYERLLRMPGGTVVLYLHGNTASRGSGHR      181
mABHD12      -------------------------GKDQMWYEDALA-SNHAIILYLHGNAGTRGGDHR      186
hABHD12      -------------------------GKDQMWYEDALA-SSHPIILYLHGNAGTRGGDHR      186
                                      ::*::**  *   .  ::******:.:**..**
```

Acyltransferase motif

```
CG15111      SEVYKLLRKLNYHVFSFDYRGYADSDPVPPTEEGVVRDAMMVFEYIAN-TTSNPIVVWGH      240
mABHD12      VELYKVLSSLGYHVVTFDYRGWGDSVGT-PSERGMTYDALHVFDWIKARSGDNPVYIWGH      245
hABHD12      VELYKVLSSLGYHVVTFDYRGWGDSVGT-PSERGMTYDALHVFDWIKARSGDNPVYIWGH      245
              *:**:* .*.***.:*****:.**  . *:*.*:. **: **::*   : .**: :***
```

Nucleophilic Motif

```
CG15111      SLGTGVATHLCAKLASLRERAPRGVILESPFTNIRDEIRMHPFAKLYKNLPWFNFTISQP      300
mABHD12      SLGTGVATNLVRRLC-ERETPPDALILESPFTNIREEAKSHPFSVIYRYFPGFDWFFLDP      304
hABHD12      SLGTGVATNLVRRLC-ERETPPDALILESPFTNIREEAKSHPFSVIYRYFPGFDWFFLDP      304
             ********:*  :*.  **  * .:**********:* : ***: :*: :* *:: : :*

CG15111      MYTNRLRFESDVHVLEFRQPIMIIHAEDDVVVPFNLGYRLYRIALDGRSRTSGPVEFHRF      360
mABHD12      ITSSGIKFANDENMKHISCPLLILHAEDDPVVPFHLGRKLYNIAAPSRSFRDFKVQFIPF      364
hABHD12      ITSSGIKFANDENVKHISCPLLILHAEDDPVVPFQLGRKLYSIAAPARSFRDFKVQFVPF      364
             : :. ::* .* :: .:  *::*:***** ****:** :** **  .**  .   *:*  *
```

Conserved Aspartate

```
CG15111      GASRKYGHKYLCRAPELPGLIQKFVENYRDAVY- 393
mABHD12      HSDLGYRHKYIYKSPELPRILREFLGKSEPERQH 398
hABHD12      HSDLGYRHKYIYKSPELPRILREFLGKSEPEHQH 398
              :.  * ***: ::**** ::::*: : .
```

Conserved Histidine

**Figure S4.** Sequence alignment of hABHD12, mABHD12 and CG15111. Sequence alignment of hABHD12, mABHD12 and CG15111 showing very high sequence homology between hABHD12 and mABHD12 (~ 94%), and high conservation between the three sequences for the acyltransferase motif (highlighted in blue), nucleophilic motif (highlighted in red), conserved aspartate (highlighted in green) and conserved histidine (highlighted in yellow), that together form the enzyme active site.
